# Multi-omic analysis reveals the genetic architecture of water-deficit stress in *Phaseolus vulgaris*

**DOI:** 10.64898/2026.02.02.702316

**Authors:** Mustafa Bulut, Regina Wendenburg, Susan Bergmann, Adilson A. Pereira Júnior, Elisa Bellucci, Elena Bitocchi, Chiara Santamaria, Laura Nanni, José G. Vallarino, Ismail Dahmani, Karin Koehl, Roberto Papa, Alisdair R. Fernie, Saleh Alseekh

## Abstract

Common bean (*Phaseolus vulgaris L.*) is one of the most important grain legumes for direct human consumption. Currently, 60% of its production is estimated to be at risk due to drought. However, the genetic basis of common bean’s drought resistance is poorly understood. To this end, we assessed the genetic architecture of drought-responsive changes in a whole genome-sequenced population of 218 common bean accessions. Using multi-omics-based trait evaluation, including lipidomics, photosynthetic and agronomic traits, followed by multi-omics genome-wide association studies (moGWAS), yielded in the detection of a myriad of moQTL for photosynthesis and yield, as well as the levels of various lipids. QTL associated with glycolipids, which are integral to photosynthesis, since they constitute the major membrane components of chloroplasts, were identified. In addition, we molecularly validated several lipid-related candidate genes via *P. vulgaris* hairy root transformation as well as transient expression in tobacco. In particular, a lipoxygenase and an allene oxide synthase were identified as explaining the variation in triacylglycerol by oxylipin production. These data provide a blueprint for multi-omics-assisted improvement of crop water stress resilience.

## Introduction

Common bean (*Phaseolus vulgaris L.*) is a major source of proteins and essential nutrients. Worldwide, it is the most consumed legume species, providing up to 36% of total daily proteins and 15% of the total daily calories (http://www.fao.org/faostat/). Wild common bean extends across a broad range from northern Mexico to northwestern Argentina ^1^. Its genetic structure is organized into three eco-geographic gene pools, shaped by population differentiation under reproductive isolation and associated with distinct adaptive patterns, influenced by geographic factors. The two main gene pools are the Mesoamerican and Andean ones, which diverged from a common ancestral wild population more than 100.000 years ago ^2^. Domestication of these two geographically separated gene pools occurred independently around 8.000 years ago ^3–5^, followed by local adaptations to changing environmental conditions, leading to distinct crop diversifications and subsequent selection of landraces. Multiple introductions from the New World, coupled with intercontinental exchanges and adaptation to new agro-ecological conditions, have disrupted the spatial isolation characteristic of the Americas. This facilitated hybridization and introgression between the Andean and Mesoamerican gene pools, generating novel genotypes and phenotypes with transgressive traits for key agronomic and adaptive characteristics ^6^. Introduced into Europe during the Columbian exchange, common bean underwent admixture between the two gene pools to adapt to European agro-environments, making it an excellent model for studying rapid genetic adaptation to novel conditions ^7^.

With ongoing climate change, driven by both natural and anthropogenic increases in atmospheric CO_2,_ drought periods are becoming more frequent and severe ^8^. Depending on the severity and the developmental stage of the crop, drought can lead to up to 95% yield losses ^9^ and it is estimated that 60% of common bean is cultivated under risk of drought stress ^10^. Multiple genetic networks inducing several morphological and biochemical adjustments are triggered upon drought ^11^. Abscisic acid (ABA)-induced stomatal closure is a prompt response to maintain the water state in plants ^12^. Further, the reduction of CO_2_ on the carboxylation site significantly reduces assimilation depending on the drought severity ^13^. Prolonged stress leads to acclimation reactions, which include osmotic adjustments ^14^, reprogramming of metabolism ^15^, activation of antioxidant systems ^16^ and an adjustment of lipid composition ^17,18^. For the latter, monogalactosyldiacylglycerol to digalactosyldiacylglycerol conversion and adjustments in fatty acid saturation enhance membrane stability and prevent structural collapse^19–21^. These changes preserve thylakoid stacking, photosystem organization, and electron transport under water deficit ^20,21^. Drought also triggers shifts in lipid metabolism toward protective lipids, including alteration of triacylglycerols, which buffer free fatty acids and prevent oxidative damage ^22^.

Harnessing nature’s diversity to dissect the genetic underpinnings of various traits in multiple crop species has been exploited using genome-wide association studies (GWAS). In *P. vulgaris*, the genetic architecture of several morphological and nutritive traits has been investigated in previous years, namely pod morphology and color characteristics ^23^, flowering time ^24^ and mineral content ^25^. By contrast, multi-environmental GWAS, such as drought versus control environments, are rare and have been conducted only for a few physiological traits, namely photosynthesis ^26,27^ and biomass accumulation ^28,29^. Nevertheless, most studies conducted lack both assessments at multi-omic layers and molecular validation of the identified QTL and further tend to focus on a single gene pool, obstructing the investigation of global patterns based on historical climate data in defined geographical locations.

Hereby, performing Genotype-by-Environment (GxE) based multi-omic GWAS (mo-GWAS) across independent experimental seasons incorporating two watering regimes, namely control and water-deficit stress, on a whole-genome sequenced (WGS) global population of 218 *P. vulgaris*, we provide a comprehensive drought-specific multi-omic data set, and assessed the genetic basis of photosynthetic, lipidomics and yield-related traits in *P. vulgaris* under drought stress. Furthermore, we identified several candidate genes involved in the stress response of *P. vulgaris*. Finally, we cloned and validated the functions of a lipoxygenase and an allene oxide synthase through hairy root assays and transient overexpression in common bean and tobacco, demonstrating their roles in oxylipin production and underscoring the importance of lipid-mediated signaling during drought stress. The multi-omics data set provides a high-resolution map to improve crop resilience.

## Results

### Natural variation of water-stress-related multi-omic changes in *P. vulgaris*

Unlocking nature’s diversity in understanding crops genetic basis to altered climates, including water scarcity, is from today’s perspective of global interest for food security. Across two experimental seasons, we performed large-scale common bean trials consisting of 218 accessions retrieved from genetic resources and evaluated them on a multi-omic level to uncover water scarcity responses in common bean.

Assessing the yield performances of these different gene pools demonstrated sensitivity to photoperiod in the Andean-2 and Mesoamerican-1 groups, as reported previously in Bellucci et al. (2023; Figure 1, B). However, under control conditions, accessions of the Mesoamerican-1 group showed partial flowering and pod setting late in the cultivation period (Figure 1, B). While holistically, the yield performances within the gene pool across both watering conditions do not differ, each genotype responded differently within the pool, with increased or reduced yield performance. The same holds true for most photosynthetic parameters. While the median levels of Φ_2_ are reasonably similar across all water-deficit-stressed and control plants, gene pool-based responses vary with differing watering volumes, as indicated by the significant changes within each treatment (Figure 1, B). To this end, we mainly emphasize the GxE covariate differentials in identifying drought-response genetic determinants in phenotypic expression. During the cultivation periods employed here, plants received over 60L and 30L of water in controlled and drought-stressed conditions, respectively (Figure 1, D). In addition, the cultivation environment was characterized by an average air temperature of 22 °C, which exceeded 40 °C on two days, and relative humidity in the range between 16% to 93%, averaging at 65% (Figure 1, C). In addition, investigating bioclimatic variables of the geographical locations from which the landraces originate indicated the highest coefficient of variations of precipitation in the Mesoamerican-1 gene pool (Figure 1, A), concurrently with the lowest precipitation in the driest month and coldest quarter. Furthermore, geolocations of this particular origin of domestication exhibit the highest mean diurnal temperature range.

**Figure 1.**
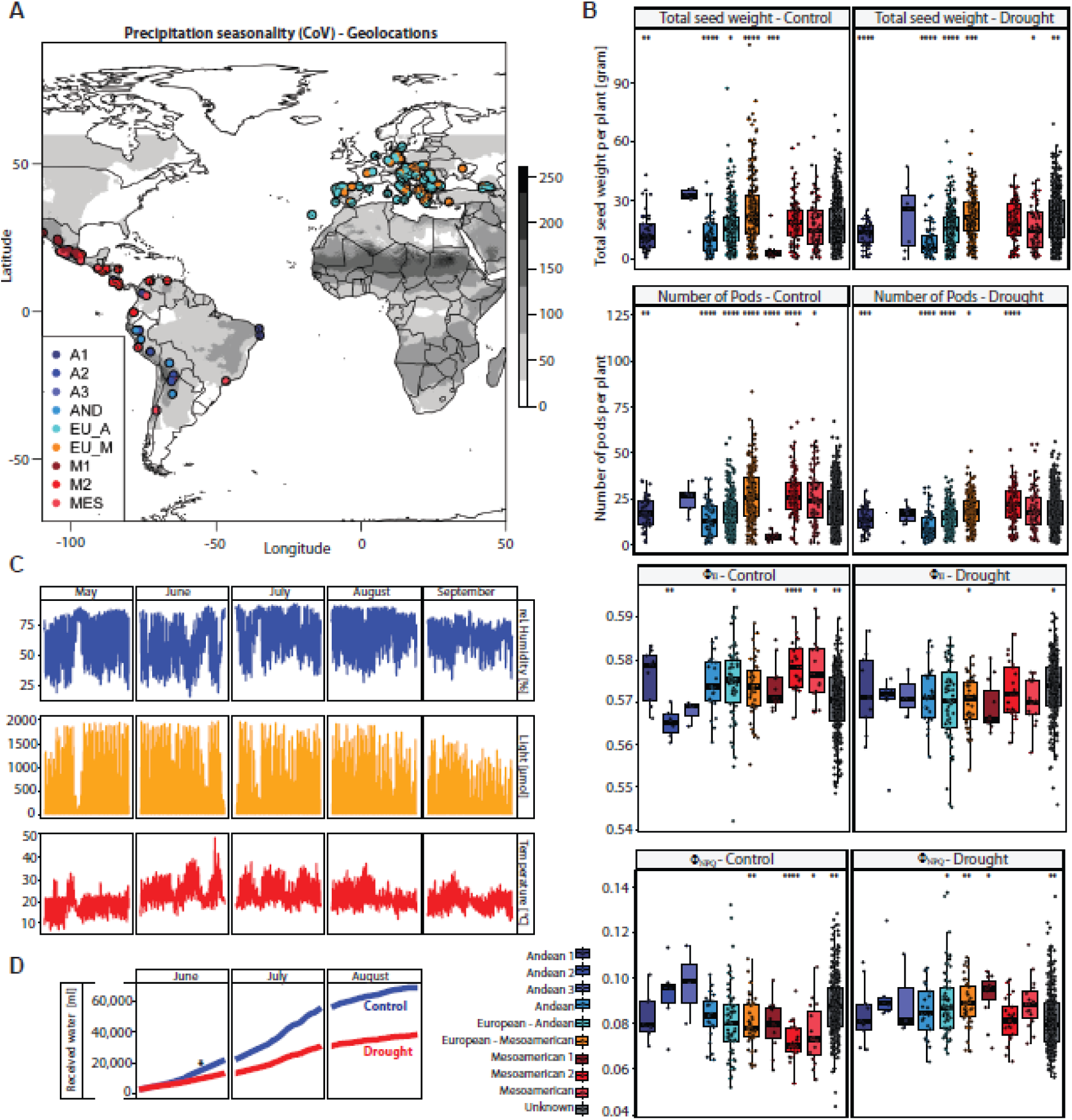
Geo-environmental and morphophysiological drought response data on the *P. vulgaris* population. **(A)** The geolocation representation of the core 218 accessions with indication of their domestication history in different colors, namely two genetic Mesoamerican subgroups (M1 and M2), three Andean genetic subgroups (A1, A2 and A3) intermediate admixed European genotypes (European Mesoamerican and Andean, depending on their proportion of the respective genetic origin) and non-associated genetic subgroups within Andean and Mesoamerican from Bellucci, et al. ^7^. Grey scale indicates the precipitation seasonality from the average across the years 1960 - 2000 calculated by the coefficient of variation (CoV) within individual annual ranges. This value represents the bioclimate variable BIO15 of the WorldClim database. **(B)** Boxplots displaying selected yield as well a photosynthetic parameters for drought and control conditions, where plants are grouped based on their domestication history in their respective group. Accessions, which were not sequenced in Bellucci, et al. ^7^, are grouped as unknown. **(C)** Exemplary environmental climate data, namely relative humidity, light intensity and temperature over the second growing season in Potsdam, Germany. **(D)** Total amount of water received by each plant across its cultivation period. Blue and red represent control and drought-stressed plants, respectively. The star indicates the time point of leaf harvest for metabolic analysis as well as for optical measurements. * = p-value <0.05, ** = p-value <0.01 and *** = p-value <0.001 of one-way ANOVA. Andean 1: *n = 87*; Andean 2: *n = 24*; Andean 3: *n = 19*; Andean: *n = 158*; European – Andean: *n = 537*; European – Mesoamerican: *n = 383*; Mesoamerican 1: *n = 53*; Mesoamerican 2: *n = 197*; Mesoamerican: *n = 147* and Unknown: *n = 1161*.

### Genetic architecture of common bean yield responses to drought

Drought has a huge impact on crop yield performance; hence, breeding of accessions that are more resilient is highly desired. The yield-related traits data obtained under water-deficit stress experiments were used to determine the covariates of the genotype-by-environment interaction in the common bean GWAS under drought and control conditions. Defined traits, namely the differential of average seed weight, resulted in a highly significant association on chromosome 6, while the differential of average pod weight displayed trends of associations towards genomic regions below the defined adjusted Bonferroni significance threshold (Figure 2, A and 3, A). Nevertheless, for the latter, a candidate gene was identified to potentially play a role in plant yield performance. When investigating the haplotype distributions for both highlighted trait-SNP associations, it became evident that the variation in the haplotypes is mainly present in the Andean gene pool, while most of the Mesoamerican accessions preserved the major allele (Figure 2, B, and 3, B). Although its distribution for average seed weight represents a minor allele (<5%), for average pod weight, the minor allele is more frequent (>5%). For both candidates, orthogroups were generated on the basis of protein sequences and explored for characterized orthologues in more studied organisms such as *A. thaliana*, resulting in the identification of *SQUAMOSA PROMOTER BINDING PROTEIN-LIKE 7* (*SPL7*) and *GPI-ANCHORED PROTEIN LORELEI 1* (*LRE*) as candidate genes for average seed weight and pod weight under drought stress, respectively (Figure 2, C and 3, C).

**Figure 2.**
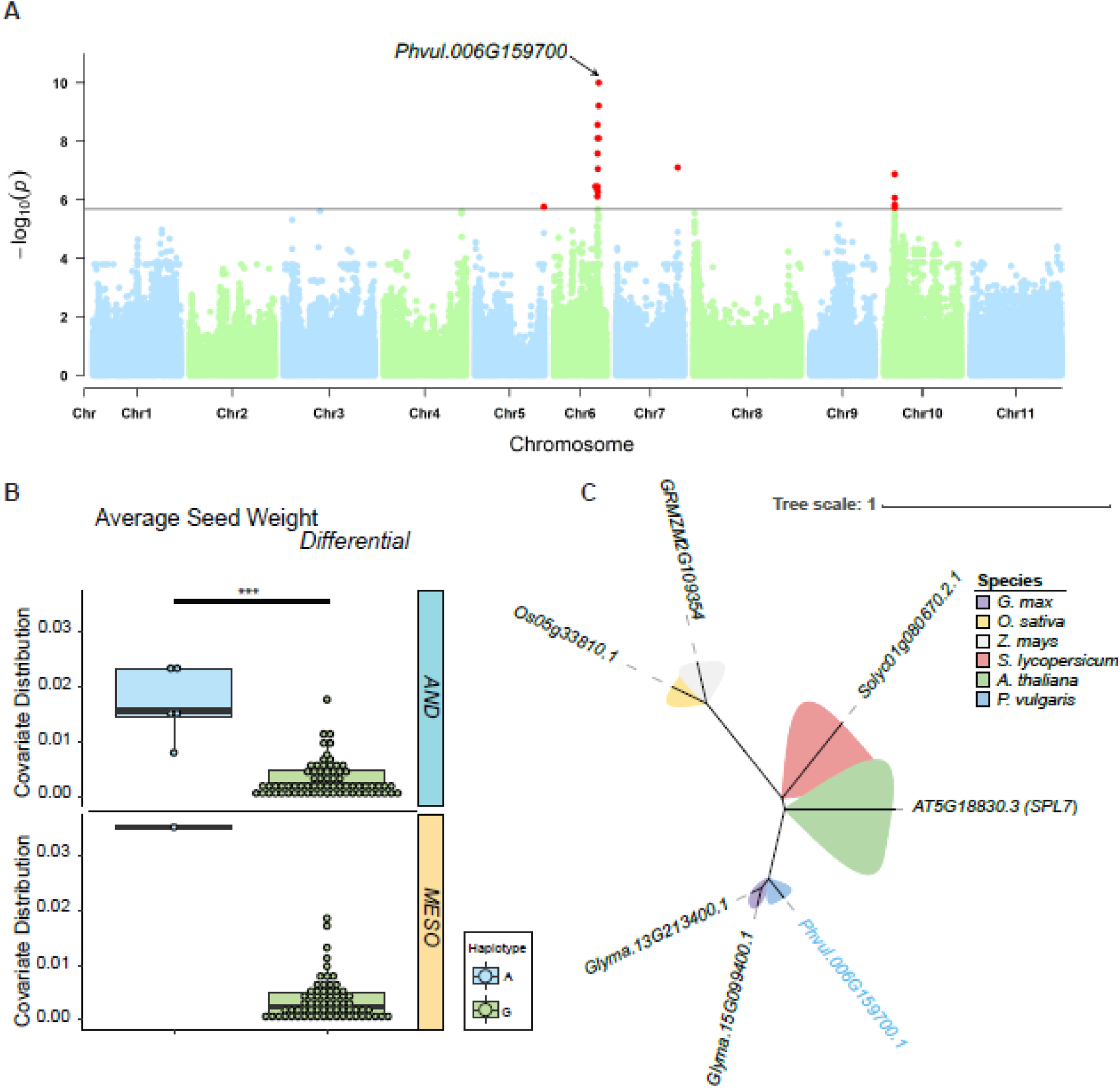
Genetic basis of drought-induced yield changes. **(A)** Genome-wide association study of average seed weight _differential_. Manhattan plot highlighting the highest association to the marker of interest with the potential candidate gene Phvul.006G159700, SQUAMOSA PROMOTER BINDING PROTEIN-LIKE 7, on chromosome 6 underlying the variation in average seed weight differential. **(B)** The boxplots represent the haplotype covariate distribution for the highlighted SNP in blue and green across both gene pools, Andean and Mesoamerican, respectively. **(C)** The unrooted phylogenetic tree reflects the orthogroup containing the gene Phvul.006G159700 highlighted in blue among orthologs across several plant species, namely Solanum lycopersicum, Arabidopsis thaliana, Phaseolus vulgaris, Glycine max, Oryza sativa and Zea mays in red, green, blue, violet, yellow and grey, respectively. *** = p-value <0.001, AND = Andean and MESO = Mesoamerican.

**Figure 3.**
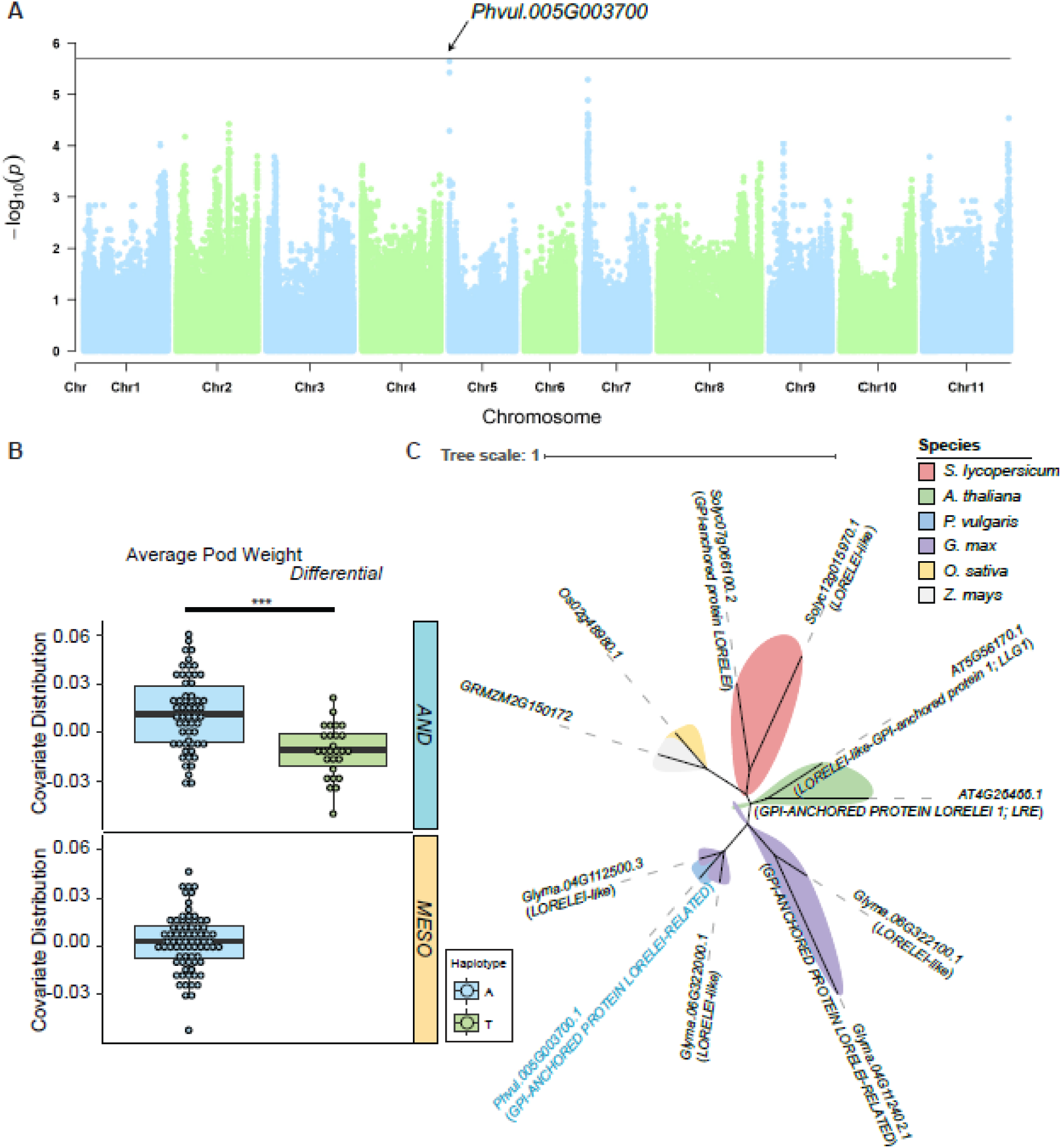
Genetic basis of drought-induced yield changes. **(A)** Genome-wide association study of average pod weight _differential_. Manhattan plot highlighting the most significant marker with the potential candidate gene, Phvul.005G003700, a GPI-ANCHORED PROTEIN LORELEI-RELATED, on chromosome 5 underlying the variation of the average pod weight differential. **(B)** The boxplots represent the haplotype covariate distribution for the highlighted SNP in blue and green across both gene pools, Andean and Mesoamerican, respectively. **(C)** The unrooted phylogenetic tree reflects the orthogroup containing the candidate gene Phvul.005G003700 highlighted in blue among orthologs across several plant species, namely Solanum lycopersicum, Arabidopsis thaliana, Phaseolus vulgaris, Glycine max, Oryza sativa and Zea mays in red, green, blue, violet, yellow and grey, respectively. *** = p-value <0.001, AND = Andean and MESO = Mesoamerican.

### Drought-responsive photosynthetic QTL discovery

Recent developments in high-throughput optical measurement units have enabled the assessment of photosynthetic parameters in larger populations, thereby facilitating the dissection of their natural variation. On-site measurements of photosynthetic activity are affected by the light conditions to which the leaves were exposed during the measurement. To circumvent such distortions, multiple measurements were taken across several days and times across the light period, yielding over 7,000 data points. By generating a linear-mixed model (LMM, see M&M), covariates of the GxE interactions and their differentials were obtained to perform GWAS. This resulted in the identification of several significant associations for several photosynthetic parameters, namely Soil Plant Analysis Development (SPAD), Φ_NPQ_as well as NPQ_t_. Among these, we highlight the GxE covariate differential-based association for Φ_NPQ_ (Figure 4, A). It became clear when dissecting the haplotype distribution for the significant SNP that observed natural diversity in those that is mainly attributed to the Andean genepool (Figure 4, B). The QTL harbors among others *Phvul.002G216900*, a gene of which the orthologues in *A. thaliana* (AT1G58440: SQUALENE EXPOXIDASE 1/DROUGHT HYPERSENSITVE 2; AT2G22830: SQUALENE EPOXIDASE 2; and AT4G37760: SQUALENE EPOXIDASE 3) have been characterized to catalyze the conversion of squalene to 2,3-oxidosqualene and further displayed drastic hypersensitivity towards drought due to an altered stomatal aperture ^30^ (Figure 4, C). Furthermore, another candidate in the defined QTL is *Phvul.002G216200,* the orthologue of AT4G04850, a K+ EFFLUX ANTIPORTER 3 (KEA3) known to be important in adaptation to fluctuating light intensities ^31–33^ (Figure 4, C). Investigating the variants for KEA3 in the *P. vulgaris* population results in four detected variants, in which one variant harbors a non-synonymous mutant in the NAD(P)-binding domain (T_634_A), while the other variants are conserved in the predicted domains. The variation within the other variants originates from C-and N-terminal non-synonymous mutants up- and downstream of the functional domains, respectively.

**Figure 4.**
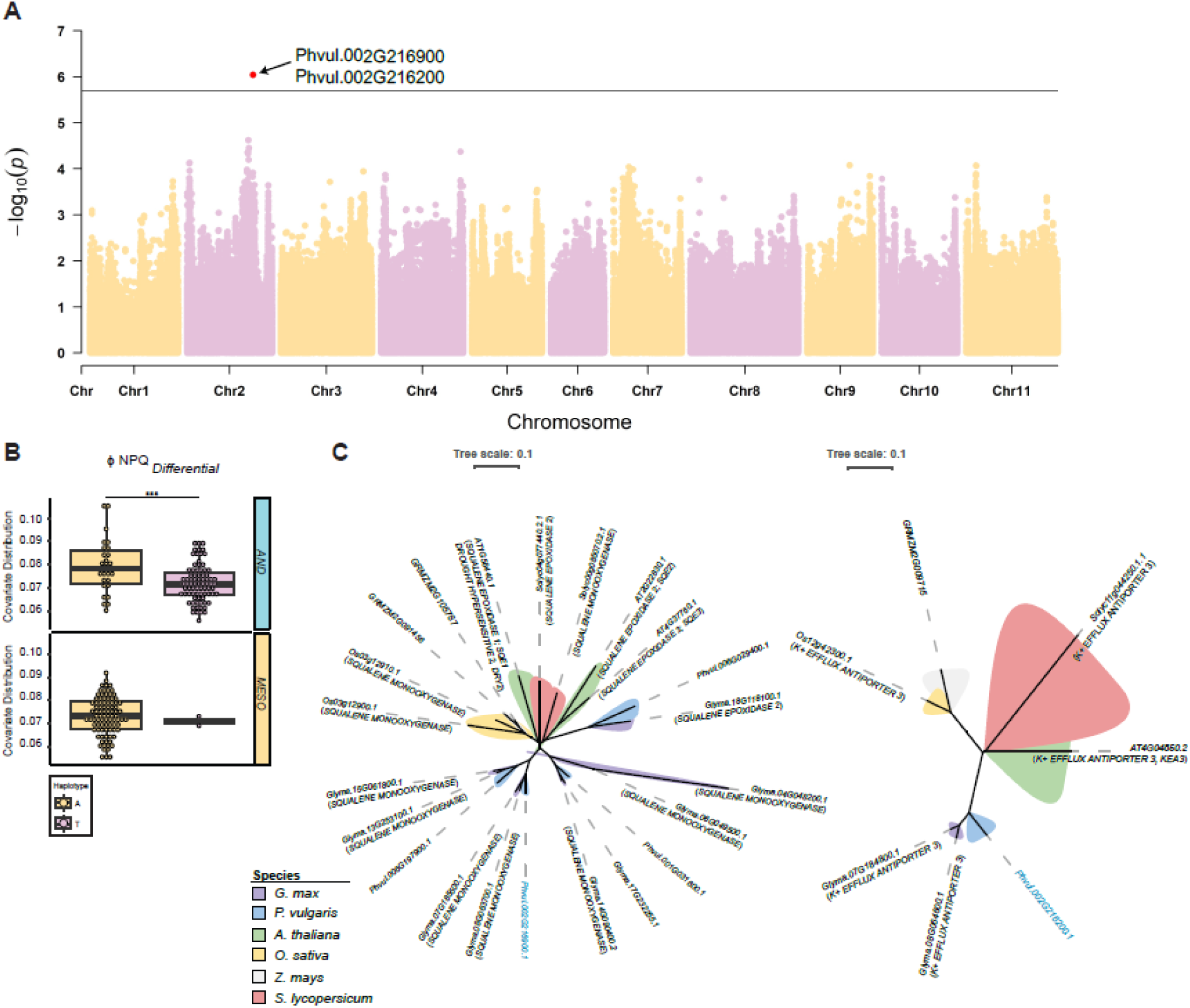
Genome-wide association study of Φ_NPQ differential_. **(A)** Manhattan plot highlighting a significantl associated SNP (red dot) and a QTL harboring potential candidate genes, namely Phvul.002G216200, with functional prediction as K+ EFFLUX ANTIPORTER 3, and Phvul.002G216900, with functional annotation as SQUALENE EXPOXIDASE 1, on chromosome 2 underlying the variation of NPQ differential. **(B)** The boxplots represent the haplotype covariate distribution for the highlighted SNP in yellow and pink across both gene pools, Andean and Mesoamerican, respectively. **(C)** The unrooted phylogenetic trees reflect the orthogroups containing the gene Phvul.002G216200 and Phvul.002G216900, respectively, highlighted in blue among orthologs across several plant species, namely Solanum lycopersicum, Arabidopsis thaliana, Phaseolus vulgaris, Glycine max, Oryza sativa and Zea mays in red, green, blue, violet, yellow and grey, respectively. *** = p-value <0.001, AND = Andean and MESO = Mesoamerican.

### Lipidomic GWAS reveals relevance of lipid signaling upon water-deficiency

Using HPLC-MS based lipidomic analysis resulted in the detection of diverse lipids belonging to several chemical classes, specifically 218 targeted lipids across 20 classes and 186 untargeted lipids. Performing supervised multivariate analysis, namely partial least squares–discriminant analysis (PLS-DA) for both experimental seasons, led to clustering of control and drought samples (Figure 5, A). Whilst a strong separation was obtained for the first experimental season, the components’ power is strongly reduced for the second experimental season, likely originating from non-controlled environmental factors in the growing seasons. Further, the loading plots for both experiments display the importance of lipids in the discrimination, with major discrimination originating from unknown lipids, while the major targeted lipids behave conservatively across both watering regimes (Figure 5, A). Unlike reported specialized metabolites, lipids do not showcase clear fingerprints on the basis of their origin of domestication ^34,35^. This, however, might originate from a strong selection of untargeted lipidomic features (n_l_ = 404).

**Figure 5.**
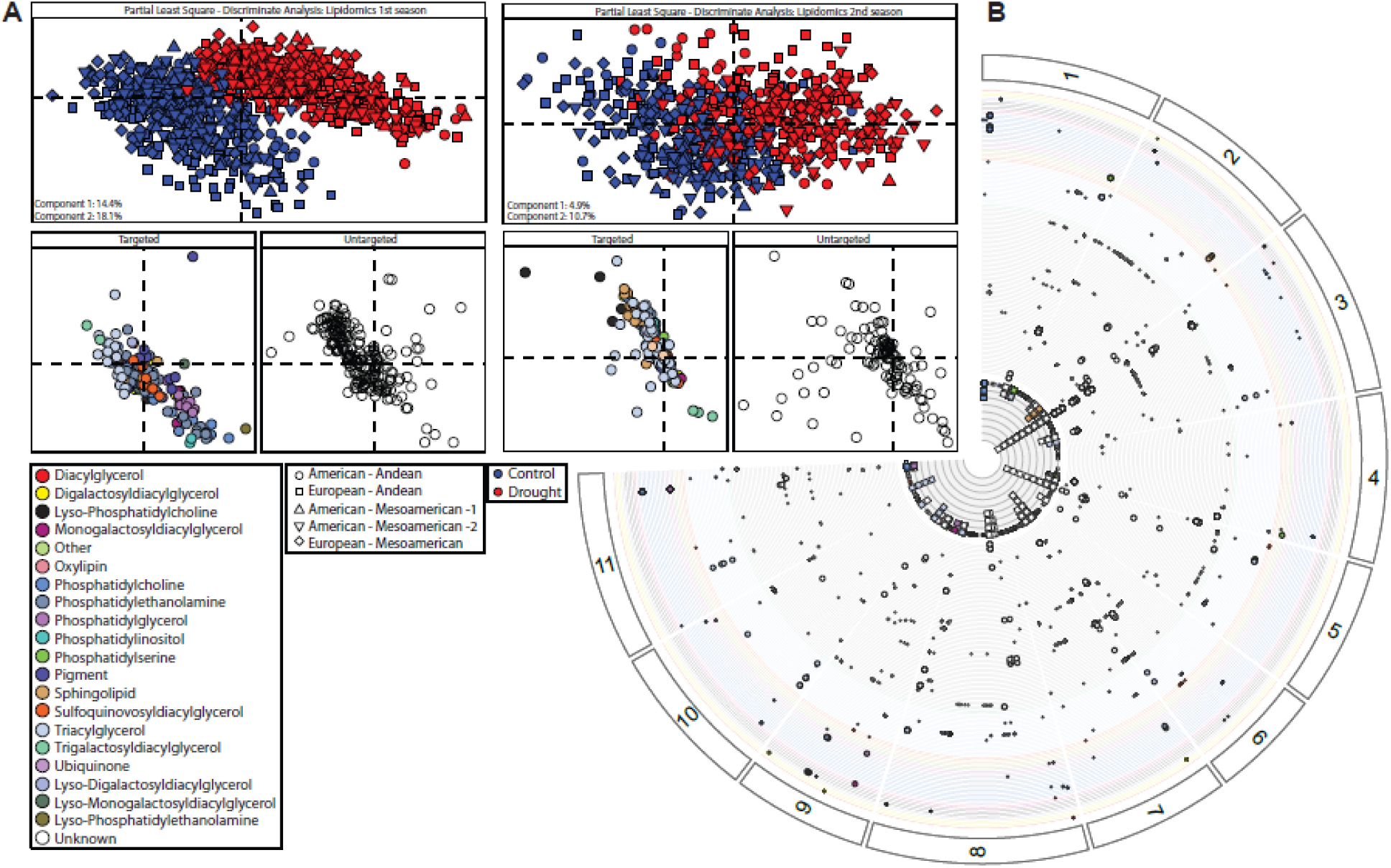
Lipidomic diversity in P. vulgaris. **(A)** 2-D projection of the partial least squares–discriminant analysis, explaining 32.2% (experimental season 1) and 15.6% (experimental season 2). Red and blu represent drought and control samples, respectively, while the shape reflects the domestication origin. Circle = ‘American - Andean’, square = ‘European - Andean’, upward triangle = ‘American – Mesoamerican -1’, downward triangle = ‘American – Mesoamerican -2’ and diamond = ‘European - Mesoamerican’. The respective loading plots of the 2-D PCA projections are separated into targeted and untargeted lipids. Colors indicate different lipid classes, while white reflects unknown lipids (untargeted). **(B)** Genetic map displaying identified QTL for targeted as well as untargeted lipids in GxE covariate differential-based GWAS (n_sig_ = 123; 72 untargeted). The colors reflect the chemical classes of each lipid. Small and big dots reflect unique and intra/inter chemical superclass-wide significant SNP associations, respectively, to highlight QTL harboring pleiotropy.

While the majority of the lipidomic GWAS resulted in similar associations between control and drought due to the strong population structure and genetic diversity, we were able to detect several unique associations for both individual genotype-by-environmental covariates and their differences across drought and control environments (Figure 5, B). To this end, several identified lipidomic QTL are highlighted with further emphasizing their importance in lipid metabolism and signaling upon environmental cues.

The first investigated candidate gene, *Phvul.003G010700,* predicted to encode an *ALLENE OXIDE SYNTHASE* (*AOS*), is obtained from environmental covariate differential based GWAS (Figure 6, A). Identification of its orthologues highlights clustering with described proteins in *S. lycopersicum* (Solyc), namely *AOS3* and *DIVINYL ETHER SYNTHASE* (*DES*), although no protein sequences from *A. thaliana* (At) were assigned to this particular orthogroup (Figure 6, B). Based on the haplotype distribution, major variation is observed in MESO accessions (Figure 6, C). *SolycAOS3* and *SolycDES* have been described in cytoplasmic catalyzation of 9*S*-hydroperoxyl-10*E*,12*Z*,15*Z*-octadecatrienoic acid (9-HPOT) downstream of *9S-LIPOXYGENASE* (*9-LOX*), yielding 9,10-EOT and colonelienic acid, respectively, while the enzymatic step catalyzed by *AtAOS1* in the generation of 12,13-epoxy octadecatrienoic acid (EOT) upstream of methyl-jasmonate (JA) biosynthesis is localized in the chloroplast. Considering the spatial separation of both pathways, tagging our protein of interest with a C-terminal *GREEN FLUORESCENCE PROTEIN* (*GFP*) suggests the involvement in the 9-LOX metabolism concluded by cytoplasmic localization (Figure 6, D). To validate our findings from the GWAS, lipidomic characterization of the constitutive *Phvul.003G010700* overexpression indeed demonstrates a significant reduction of TAG. Concurrent with this reduction, higher levels of oxylipins were detected, highlighting increased rates of their production in *P. vulgaris* hairy roots (Figure 6, E).

**Figure 6.**
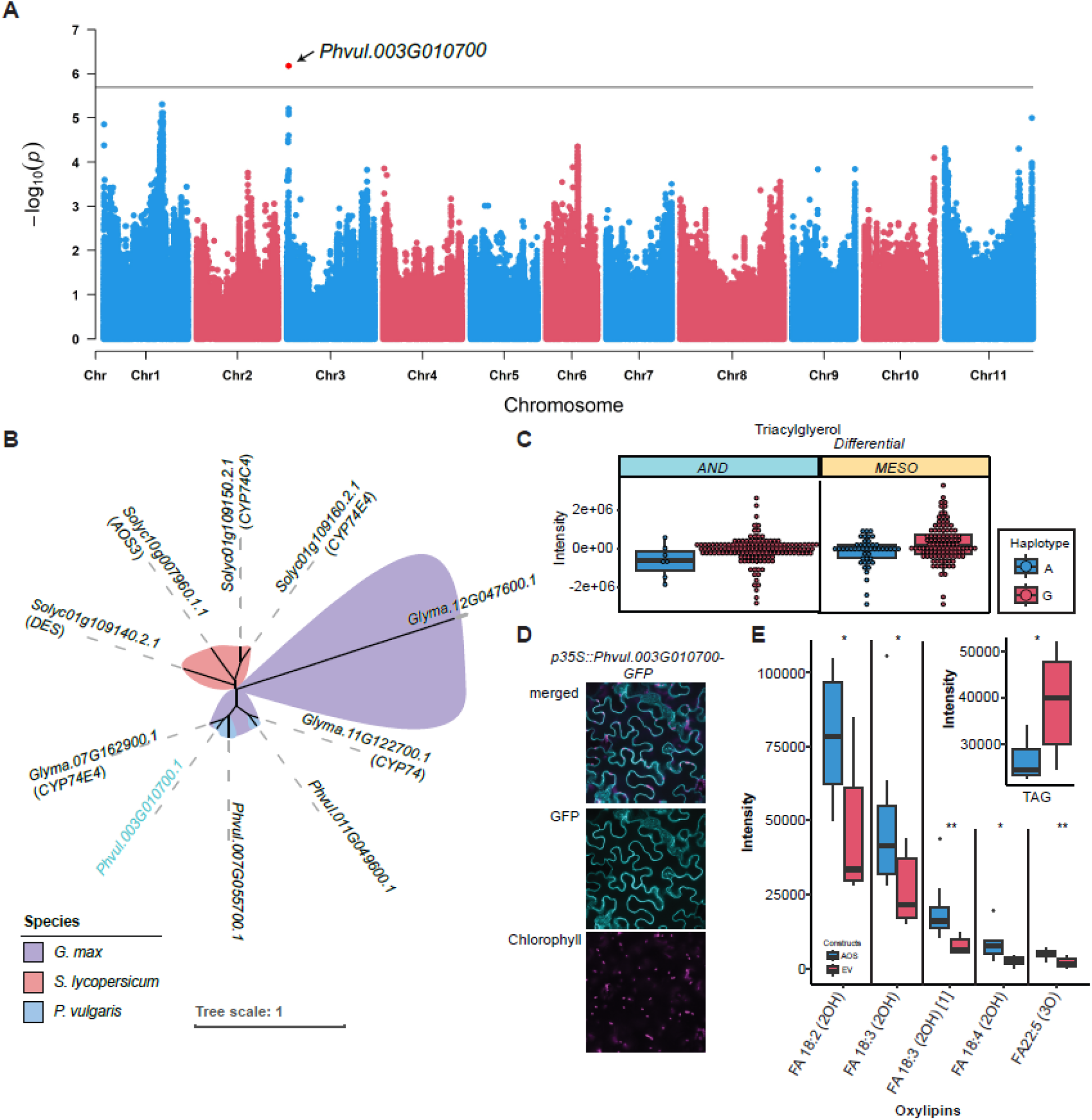
Genome-wide association study of Triacylglycerol_differential_. **(A)** Manhattan plot highlighting the significant associated marker with potential candidate gene Phvul.003G010700, functionally predicted a an ALLENE OXIDE SYNTHASE, underlying the variation of TAG. **(B)** The unrooted phylogenetic tree reflects the orthogroup containing Phvul.003G010700 highlighted in blue among orthologs across several plant species, namely Solanum lycopersicum, Glycine max and Phaseolus vulgaris in red, violet and blue, respectively. Other species included in the analysis, such as Oryza sativa, Zea mays and Arabidopsis thaliana do not have predicted orthologs within this particular orthogroup. **(C)** The boxplots represent th haplotype distribution for the highlighted SNP in blue and red across both gene pools, Andean and Mesoamerican, respectively. **(D)** C-terminal fused GFP to Phvul.003G010700, highlighting subcellular localization. Different channels for GFP and the chlorophyll autofluorescence are depicted, including the merged channels. Turquoise = GFP and magenta = chlorophyll autofluorescence. **(E)** Lipidomic analysi of transient overexpressing Phvul.003G010700 using hairy root transformation in P. vulgaris. Red and blue boxplots reflect empty vector (EV) and overexpression of the AOS, Phvul.003G010700, respectively. Overexpression of Phvul.003G010700 results in higher levels of oxygenated fatty acids (FA) and reduced TAG. * = p-value <0.05, *** = p-value < 0.01 and *** = p-value <0.001. TAG = triacylglycerol, AOS = ALLENE OXIDE SYNTHASE, GFP = Green fluorescence protein, DES = DIVINYL ETHER SYNTHASE, AND = Andean and MESO = Mesoamerican.

The second gene of interest that stood out in the GWAS, *Phvul.005G156900*, is a putative *LINOLEATE 9S-LIPOXYGENASE* (Figure 7, A), which acts upstream of the previously described candidate *Phvul.003G010700*. The orthogroup highlights two proteins for both *P. vulgaris* and *G. max,* while the proteomes of the other included species do not contain any orthologous proteins (Figure 7, B). In addition, the haplotype distribution showed major discrimination between both haplotypes based on domestication origin, namely the haplotype A and C for Andes and Mesoamerica, respectively (Figure 7, C). However, the minor allele in the Andean population displayed strong induction in contrast to the major allele upon drought. In order to identify its subcellular localization, *Phvul.005G156900* was cloned, tagged with a C-terminal *GFP* under a 35S promoter and overexpressed in *N. benthamiana*, resulting in endoplasmic reticulum (ER) and cytoplasmic localization (Figure 7, D). Furthermore, lipidomic profiling of transient overexpression in *N. benthamina* leaves and hairy roots in *P. vulgaris* highlights induction of several putative HPOTs (309.2072*m/z*) and EOTs/ colnelenic acids (291.1966*m/z*), indicating enzyme unspecificity in the generation of 9-*S* and 13-*S* HPOTs and their downstream products (Figure 7, E).

**Figure 7.**
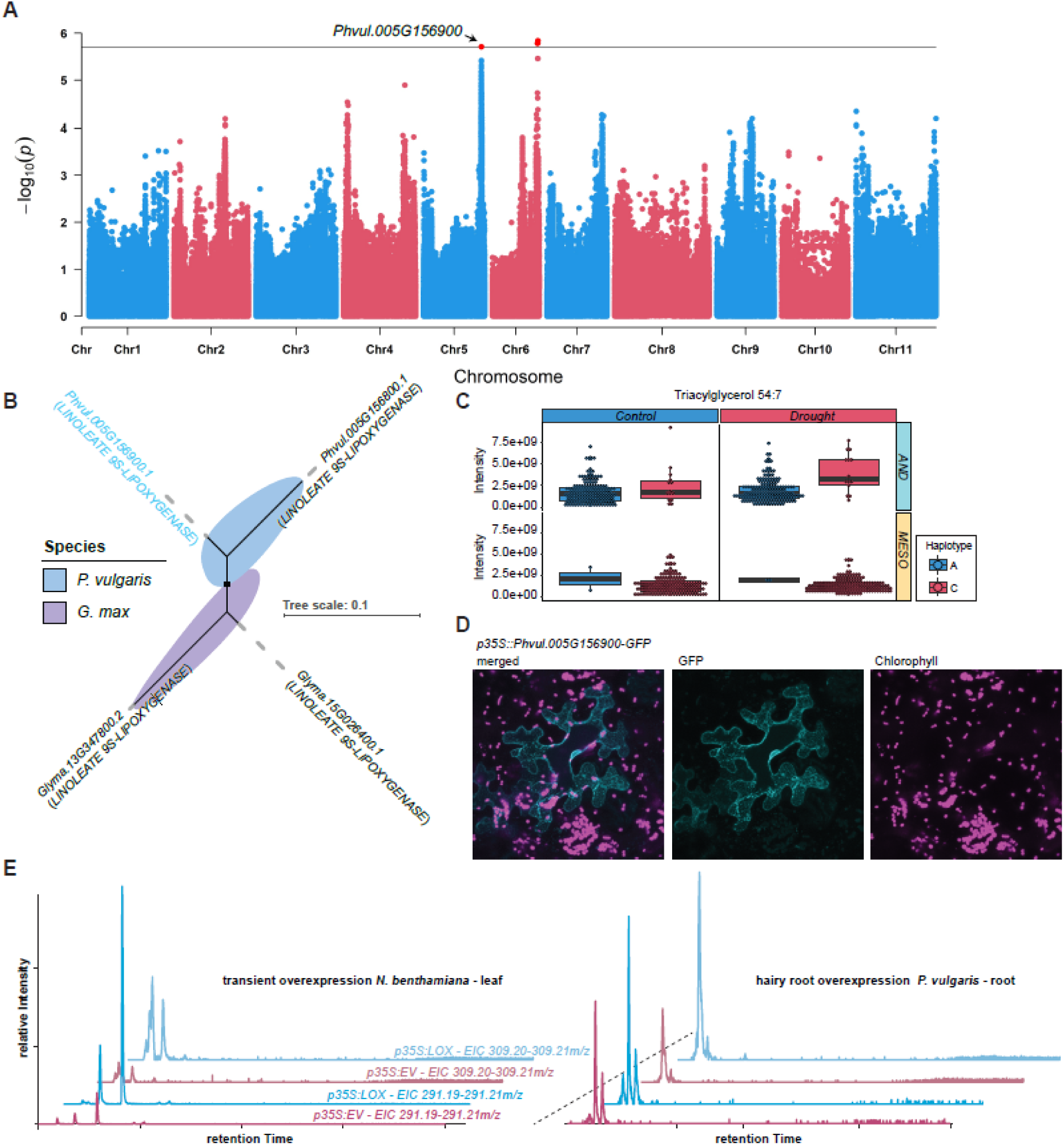
Genome-wide association study of Triacylglycerol_drought_. **(A)** Manhattan plot highlighting the significant associated marker with the candidate gene, Phvul.005G156900, with functional annotation as LINOLEATE 9S-LIPOXYGENASE, underlying the variation between TAG under control and drought conditions. **(B)** The unrooted phylogenetic tree reflects the orthogroup containing Phvul.005G156900 highlighted in blue among other orthologs across Glycine max and Phaseolus vulgaris in violet and blue, respectively. Other species included in the analysis, such as Solanum lycopersicum, Oryza sativa, Zea mays and Arabidopsis thaliana do not have predicted orthologs within this particular orthogroup, highlighting divergence in Leguminosae. **(C)** The boxplots represent the haplotype distribution for th highlighted SNP in blue and red across both gene pools, Andean and Mesoamerican, respectively, as well as control and drought treatments. **(D)** C-terminal fused GFP to Phvul.005G156900, highlighting subcellular localization. Different channels for GFP and the chlorophyll autofluorescence are depicted, including the merged channels. Turquoise = GFP and magenta = chlorophyll autofluorescence. **(E)** Extracted ion chromatogram section of hairy root as well as transient transformed leaves with empty vector (EV) in red and overexpression (35S::LOX) constructs in blue, highlighting an increase of putative HPOTs (309.2072m/z) and EOTs/ colnelenic acids (291.1966m/z). GFP = Green fluorescence protein, LOX = LIPOXYGENASE, HPOT = 9S-hydroperoxyl-10E,12Z,15Z-octadecatrienoic acid, EOT = 12,13-epoxy octadecatrienoic acid, AND = Andean and MESO = Mesoamerican.

The final lipidomic candidate investigated was a putative DIACYLGLYCEROL KINASE (DGK) obtained from DGDG 38:5 under drought GWAS (Figure 8, A). The orthogroup of this particular candidate highlights, among others, a characterized protein in *A. thaliana* (DGK2; AT5G63770) (Figure 8, B). In addition, haplotype distribution is showing allelic diversity uniquely within the Andean population, in which accessions with the G haplotype have higher levels of DGDG 38:5 under drought conditions when compared to control watering (Figure 8, C). When trying to determine the subcellular localization, aggregation of the GFP signal was observed (Figure 8, D). To narrow down potential targeted organelles, we co-infiltrated our protein of interest with a peroxisomal marker tagged with mCherry (Figure 8, D). The fact that the observed signals did not co-localize indicates accumulation of the GFP signal in a cellular compartment excluding cytoplasm, ER, chloroplast, peroxisome and nuclei. Furthermore, lipidomic profiles of transiently overexpressed DGK in *N. benthamiana* leaves highlighted a plethora of changes in both negative as well as positive ionization modes, including TAGs, PEs, PCs, lyso-PC, fatty acids and oxylipins in *N. benthamiana* leaves (Figure 8, E). By contrast, *P. vulgaris* hairy roots overexpression resulted in mainly a reduction of several lipids, highlighting species and/or tissue-specific divergence in its role and functions, as clearly depicted in the volcano plots (Figure 8, E).

**Figure 8.**
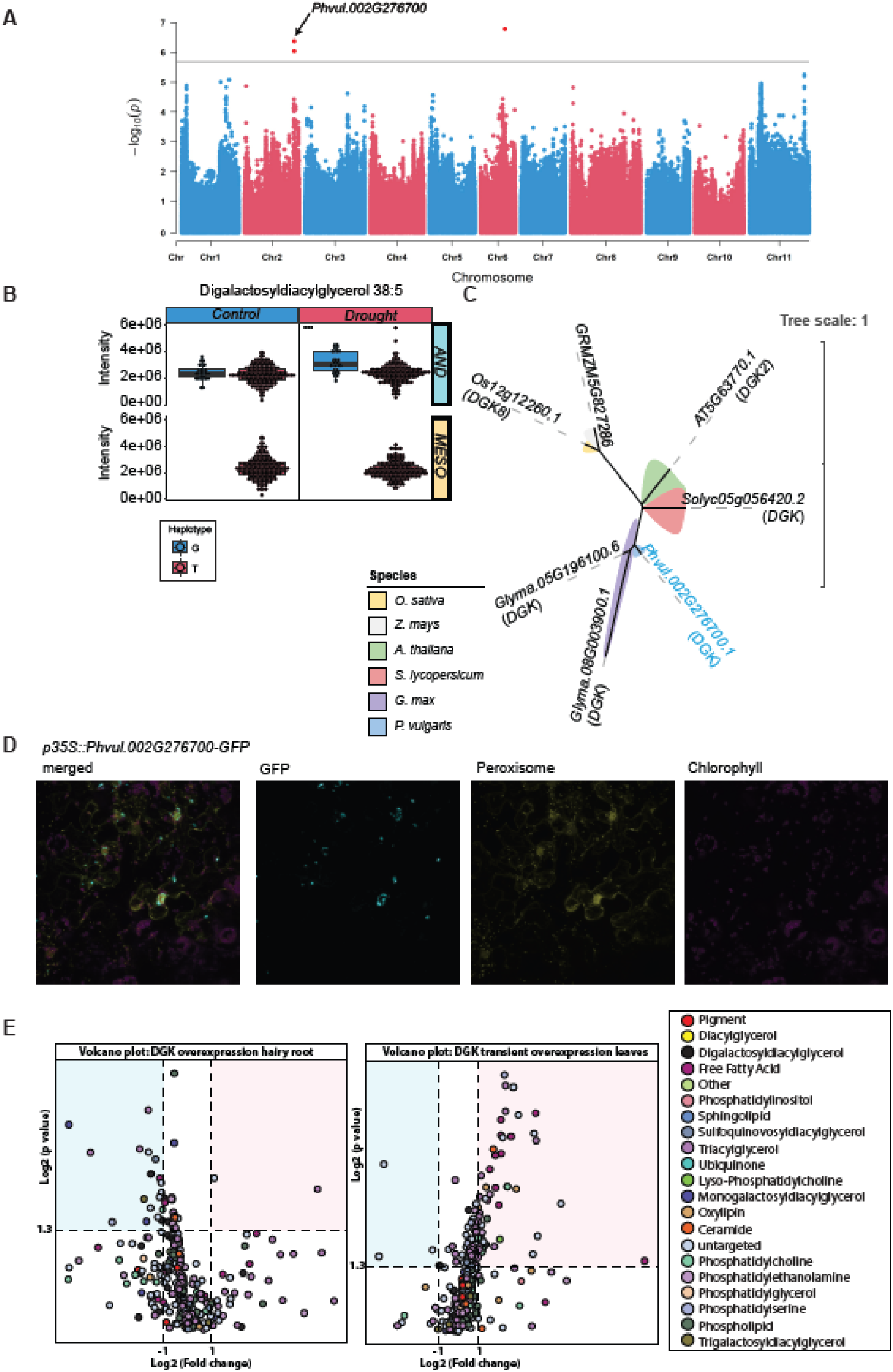
Genome-wide association study of Digalactosyldiacylglycerol_drought_. **(A)** Manhattan plot highlighting the significant associated marker with the candidate gene *Phvul.002G276700*, a functionally annotated DIACYLGLYCEROL KINASE, underlying the variation in DGDG 38:5. **(B)** The unrooted phylogenetic tree reflects the orthogroup containing the gene *Phvul.002G276700* highlighted in blue among orthologs across several plant species, namely *Solanum lycopersicum*, *Arabidopsis thaliana*, *Phaseolus vulgaris*, *Glycine max*, *Oryza sativa* and *Zea mays* in red, green, blue, violet, yellow and grey, respectively. **(C)** The boxplots represent the haplotype distribution for the highlighted SNP in red and blue across both gene pools, Andean and Mesoamerican, respectively, as well as treatments for DGDG 38:5. **(D)** C-terminal fused *GFP* to *Phvul.002G276700* transformation in *N. benthamiana* leaves, highlighting subcellular localization. Different channels for mCherry, GFP and the chlorophyll autofluorescence are depicted, including the merged channels. Yellow = peroxisomal marker (mCherry), turquoise = GFP and magenta = chlorophyll autofluorescence. **(E)** Volcano plots display significant (p-value < 0.05) log_2_(fold changes) (>1 and <-1) across hairy root (left) as well as transient Nicotiana leaf (right) transformations against their EV controls. Each dot represents a lipid, while the colors highlight the associated lipid class. The blue and red underscored area highlight significant down- and upregulated lipids. DGDG: Digalactosyldiacylglycerol, *** = p-value <0.001, AND = Andean and MESO = Mesoamerican.

## Discussion

### Yield-associated candidate genes under drought stress

SQUAMOSA PROMOTER-BINDING PROTEIN (SBP) -LIKE (SPL) proteins are transcription factors defined by a highly conserved 76 amino acid sequence called the SBP domain ^36^, with essential roles in phase separation, flower and fruit transition, as well as signaling of gibberellins and response to copper ^37–39^. While *A. thaliana* contains 17 *SPL* genes, *P. vulgaris* has 23 predicted SBP DNA-binding domain containing genes. The majority of SPLs are essential for plant development with miRNA interplay in regulating floral transition and belong to the SPL subfamily II (SPL2-6, SPL8-11, SPL13 and SPL15), while SPL subfamily I proteins (SPL1, 7, 12, 14 and 16) are less characterized yet have be reported in thermotolerance and copper and zinc deficiency ^40,41^.

From a natural variation perspective, subfamily II SPLs have been highlighted in monocot GWAS, namely SPL3 in *Hordeum vulgare* and SPL13 in *Oryza sativa* for coleoptile and grain size, respectively ^42,43^. Genome-wide analysis of the SPL gene family in *Medicago sativa* resulted in the identification of ABA-responsive elements (ABREs) in the predicted *cis*-regulatory elements. Furthermore, three SPLs, namely *MsSPL17*, *MsSPL23* and *MsSPL36*, have been reported to be upregulated upon salt stress ^44^, highlighting, next to the developmental importance, potential role in abiotic stress response.

The next drought-induced yield-related candidate gene identified, namely LORELEI-like Glycosylphosphatidylinositol (GPI) - anchored protein1 (LLG1), has been reported to be a pivotal interactor with the receptor kinase FERONIA (FER), a multifunctional regulator of plant growth and development ^45^. Besides FER, LLG also interacts with RAPID ALKALINIZATION FACTOR (RALF), a peptide hormone ^46^, building a FER-LLG-RALF protein complex involved in multiple processes in growth, immunity and stress response ^45,47,48^. Overall, *llg1* mutant exhibits retarded growth with accumulating higher anthocyanin levels, reduced rosette size, decreased sensitivity to auxin-stimulated ROS production and sensitivity to abscisic acid ^45^. Furthermore, FER was shown to recover root development after an ionic stress, namely high levels of salt, by eliciting cell-specific calcium transients to maintain cell-wall integrity ^49^. While the orthologue of LLG1–LRE is important for reproductive growth as mutants display delayed seed development and abortion, *llg1* mutants are fully fertile and do not show additive defects in *lre* pistils ^50^. This suggested that LLG1 and LRE are non-redundant proteins in pollen development clustering in the same orthogroup, whereas the proteome of *P. vulgaris* contains one assigned protein, highlighting potential LLG1 and LRE bi-functionality in *P. vulgaris*. Overall, to the best of our knowledge, this is the first GWAS identifying subfamily I SPL and LLG in drought-induced yield performance changes.

### Photosynthetic variation

The *dry2/sqe1* mutant in *Arabidopsis* is characterized by hypersensitivity to drought, altered stomatal response and root deficiency ^30^. SQE1 catalyzes the conversion of squalene to 2,3-oxidosqualene upstream of the triterpene pathway, which yields, among others, brassinosteroids. These mutants showed differences in root-specific steroidal accumulation, with changes in reactive oxygen species originating from stomatal and root defects ^30^. From a biotic stress-physiological perspective, legumes such as common bean rely on the production of triterpene saponins downstream of 2,3-oxidosqualene as a result of herbivory, which, by contrast, is not the case in *Arabidopsis* due to the evolutionary divergence of other defense compound classes such as glycosinolates in Brassicaceae ^51^. Thereby, hypothesizing on molecular mechanisms based on the basis of *Arabidopsis* studies can be arbitrary. Nonetheless, presuming similar mechanisms in *P. vulgaris* germplasm, influence on stomatal aperture can result in a plethora of downstream alterations, such as changes in radical formation and in excited energy release via non-photochemical quenching. The latter of which would inherently lead to an increase in the proportion of photonic energy towards NPQ, while simultaneously reducing that towards Φ_II_.

On the other hand, tight regulation of lumen acidification in maintaining photosynthetic integrity is crucial under fluctuating environments. K+ EFFLUX ANTIPORTER 3, an antiporter localized in the thylakoid membrane, was shown to be critical for high photosynthetic efficiency, as mutants of this antiporter exhibit prolonged dissipation of absorbed light energy as heat, namely induction of non-photochemical quenching, in the transition of high light to low light as well as light induction from dark to low light conditions ^31^. As for drought, it was shown that common bean shows a reduction in ATPase abundance ^52^. To this end, the de-acidification of the lumen would depend on bypasses in releasing protons to the stroma to present an energy imbalance, such a bypass could potentially be provided by KEA3. Moreover, the lumen-localized C-terminal end of KEA3 is attributed to harbor regulatory functions ^53^. Deletion of the C-terminal end results in reduced feedback-regulation and increased photodamage during prolonged light stress, while carbon fixation is increased in the initial photosynthetic induction in high light ^54^. PvKEA3 consists of two major domains, the cation/H^+^ exchanger (AA_116-503_) and the NAD(P)-binding domain (AA_534-674_). By investigating the C-terminal variation within the NAD(P)-binding domain of the *P. vulgaris* germplasm, we could demonstrate one distinct amino acid variation in T_634_A. In humans, a change in T_188_A in a NAD^+^-dependent 15-hydroxyprostagladin dehydrogenase led to inactivation and demonstrated the essentiality of T_188_ in NAD^+^ interaction and enzyme activity ^55^, while a cytochrome P450 underwent rapid autoinactivation in T_303_A when electrons were provided by NADPH ^56^. Assuming KEA3 inactivation due to lack of NAD(P) binding in accessions possessing the A_634_ would potentially support the observed differences in Φ_NPQ_. Diving into the SNP variation leading to the amino acid change clarifies the presence of A_634_ in American–Mesoamerican–2 and some European–Mesoamerican accessions. By contrast, considering the overall relatively minor changes observed in Φ_NPQ_ as depicted in Figure 1, the variation might originate from the differing light intensities plants were acclimated to, which would be in line with all previously described functions of KEA3. Furthermore, in regards to fluctuating light conditions, the thylakoid ion transport is influenced by light-environmental factors and has a profound effect on xanthophyll portioning, photoprotection, photosynthetic efficiency and lumen pH in the modulation of dynamic photosynthesis ^33^. Ultimately, either presumed mode of action of the candidate genes will require considerable further investigation.

### Involvement of lipid signaling under drought

The role of lipid signaling in response to biotic, as well as abiotic, stress is pivotal and well documented in the literature ^57,58^. Strikingly, to this end, we identified two QTL harboring genes involved in consequent enzymatic steps of oxylipin biosynthesis, namely the hydroperoxydation of fatty acids (*Phvul.005G156900*) followed by the rearrangement of the hydroperoxides (*Phvul.003G010700*). The oxylipin pathway is well established, especially the biosynthesis of jasmonates, for which these particular enzymatic steps are localized in plastids. In our co-localization experiments, we could prove an ER/cytoplasmic localization for both of our proteins of interest, suggesting that they do not participate in jasmonate biosynthesis. Furthermore, we showed that the overexpression leads to induction of (i) several HPOTs (309.207*m/z*) and EOTs/colnelenic acid (291.1966*m/z*) as shown in Figure 7E, while (ii) simultaneously inducing downstream oxylipin production (Figure 6, E). Based on the induction observation of several oxidized fatty acids, we hypothesize that product unspecific catalyzation, namely by 9*S*-LOX and AOS activity, occurs. This dual activity would differ from the traditional LOX activities described *in A. thaliana* ^59^, however, in other plant species, such as maize, the dual conversion of α-linolenic acid into 9- and 13-hydroperoxylinolenic acids is more commonly reported ^60^. While 12-OPDA is well investigated, the activity of 10-oxo-11-phytoenoic acid (10-OPEA), derived from 9-LOX, is less studied. A report highlights the production of so-called ‘death acids’, including 10-OPEA and related cyclopente(a)nones, and their function as cytotoxic phytoalexins as well as transcriptional mediators upon *Cochliobolus heterostrophus* infestation in maize ^61^. Further, several studies highlight the involvement of lipoxygenases in abiotic stress responses – particularly ionic as well as oxidative stress ^62,63^. Recently, Lung, et al. ^64^ demonstrated ACYL-COA-BINDING PROTEIN3 (ACBP3) and ACBP4 as negative regulators of the vegetative lipoxygenase VLXB by means of sequestering it at the ER in a linoleoyl/linolenoyl-CoA–dependent manner. Moreover, salt-stress-induced alternative splicing disrupts this interaction, enhancing LOX activity and salt tolerance in *G. max*. Interestingly, the orthologous genes in *G. max*, which were reported for the presented LOX (orthogroup: *Phvul.005G156900*, *Phvul.005G156800*, *Glyma.15G026400*, *Glyma.13G347800*), were both regulated by the GmACBP4.1. Given that salt-stress is an osmotic stress, which is also prevailing under water-deficiency, a similar mechanism can be assumed in the potential regulation of this LOX. Oxylipins are highly bioactive; for example, their functional convergence with ABA in the control of stomatal apertures has been reported ^65^ and highlights the production of 12-OPDA and not JA upon drought, indicating an uncoupling of JA production. However, based on the obtained phenomics data, including ambient and leaf temperature, which can serve as a proxy for stomatal aperture, no significant differences could be observed for the plants when grouped on the haplotypes, resulting in the oxylipin variation. Furthermore, plants in which 12-OPDA was induced were more drought-tolerant due to exhibiting reduced stomatal apertures ^65^. Intriguingly, exogenous treatment with 12-OPDA has likewise resulted in the promotion of stomatal closure in *Solanum lycopersicum* and *Brassica napus* ^65^. Previous GWAS conducted on other plant species, namely *O. sativa,* highlighted, among others the identification of LOX as being important for early vegetative growth under salinity^66^. However, the current study is the first of its kind in identifying candidates of oxylipin biosynthesis involved in lipidomic changes upon drought. While LOX in itself was identified in GWAS studies and has been reported to be a major contributor in flavor volatile production in tomato fruits ^67^, no study has been reported to date that identified a lipidomic QTL harboring natural variation in a *CYP74*.

Lastly, autophagy is known to be a sophisticated regulator of plant growth and its interplay with environmental signals. Several autophagy-related genes were identified as playing a major role in the regulation of drought resistance ^68^. Based on the observed GFP signal, the assumption of DGK2 localization in autophagosomes was made. DIACYLGLYCEROL KINASE catalyzes phosphorylation of diacylglycerols (DGs), yielding phosphatidic acid (PA). In mammalian cells, PA originating from phospholipase D (PLD) and DGK can regulate mTORC1 activity. Further, inhibition of DGK was shown to induce autophagy and apoptosis ^69^. This would contradict our results, where overexpression seems to induce autophagy. However, trafficking of DGK into the autophagosome would not only indicate the presence of ongoing autophagy, but also indicate recycling of the DGK protein itself. This might be an autoprotective mechanism in the degradation of DGK to stop TORC induction based on feedback regulation. Further, induction and reduction of several lipids are observed for leaves and roots, respectively. However, untangling these observations would need considerable further analysis.

### Natural variation across domestication origins and cross-omic comparison

Using the GxE covariate (differential) based moGWAS allowed the detection of a myriad of novel mo-QTL and the dissection of drought/water scarcity inductive regions of the *P. vulgaris* genome. Interestingly, contrary to expectations, the most significant SNPs identified for the highlighted traits exhibit substantial haplotype variation in the AND population, particularly for the morphophysiological traits. In contrast, the MESO subpopulation displays higher polymorphic variation across the genome. Although lipid metabolism and its remodeling are well documented as essential for maintaining chloroplast integrity under osmotic and oxidative stress^21^, we did not detect clear associations between lipid profiles, particularly galactolipids, and photosynthetic parameters such as LEF, Φ2, NPQ, or yield-related traits. Similarly, comparison of lipidomic and phenomic QTL revealed no substantial co-localization, likely reflecting the higher complexity and polygenic nature of agronomic and phenomic traits relative to (un)targeted lipid traits. Nevertheless, multi-omic approaches remain crucial for resolving the mechanistic links among lipid metabolism, photosynthesis, and yield to better understand plant resilience. As noted above, leaf temperature could not be reliably used to test oxylipin-mediated stomatal regulation; pronounced diurnal variability in temperature measurements prevented consistent interpretation. Nonetheless, the collective results concerning water-stress-induced changes on multi-omic layers in *P. vulgaris* indicate that the genetic regions underlying these changes can help to improve plants’ resilience to drought by incorporating this knowledge into multi-omics assisted crop breeding.

## Methods

### Genetic resource

The genetic resource of the core population was described previously in Bellucci, et al. ^7^. In brief, it consists of 199 domesticated whole genome sequenced (WGS) single seed descent (SSD) and an additional 19 seeds directly sampled from the providing seedbank originating from the Americas and Europe with approximately 3.6 million SNPs. Raw sequencing data used in this study are available in the NCBI Sequence Read Archive under BioProject PRJNA573595. DOIs for the set of analysed accessions are listed in Supplementary Data 1 of Bellucci, et al. ^7^. With respect to its domestication, the population is equally distributed between Mesoamerican and Andean. The genetic material was provided by UNIVPM in the framework of the European project INCREASE (https://www.pulsesincrease.eu/). Further, the bioclimatic variables of their geographical locations were retrieved from the WorldClim database using *raster* functions in R.

### Experimental design and phenotyping

Plants were grown in two independent seasons, the summer of 2019 and 2021, in semi-field conditions of the Max-Planck Institute of Molecular Plant Physiology (Golm, Germany). Each genotype was grown in triplicate and duplicate in 2019 and 2021, respectively. Two distinct water regimes were introduced, control and drought, in which drought was defined as receiving 50% of the amount of water control plants received, ending up with a dozen of hundred plants for either experimental season, respectively. Plants were sown in early May and transferred to pots. Stress application started in June. After 10 days of water treatment, leaves were harvested for metabolic characterization. Throughout the cultivation period, phenotypic traits, such as growth habit, flowering time and flower color, were assessed. Finalizing the experiments, yield of each plant was determined during September. More information on the accessions can be found in Bellucci, et al. ^7^. Climate data during the cultivation time is provided in Figure 1C.

### Photosynthetic measurements

Optical measurements were taken starting ten days after the onset of drought across consecutive days on a side leaflet of the third fully developed leaf using MultispeQ 2.0 devices calibrated with CaliQ calibration system ^70^. The light protocol ‘Photosynthesis Rides 2.0’ was used. In brief, this protocol measures ambient PAR and simulates it before taking optical measurements. The data collection was conducted during the morning, 3h after dawn. Data cleaning and refitting of optical parameters, namely ECStau and g_H+_, were performed over a linear regression ^71^. Measurements include total magnitude of electrochromic shift (*ECSt mAU*), rate of proton flux (*g_H+_*), linear electron flow (*LEF*), estimated non-photochemical quenching (*NPQ_t_*), quantum yield of photosystem II (Φ*_2_*), ratio of incoming light that is lost via non-regulated processes (Φ*_NO_*), ratio of incoming light that goes towards NPQ (Φ*_NPQ_*), proton conductivity (*v_H+_*) and relative chlorophyll content (SPAD). The raw data points can be assessed under the photosynQ.org project ID ‘7263’ and ‘12397’ for the experiment in 2019 and 2021, respectively. For the first season, one data point was generated for each plant, while for the second experiment, four data points were obtained. For the GWAS, a mixed linear model was generated explaining each optical parameter (y_o_) as follows for the data point of project ‘12397’:

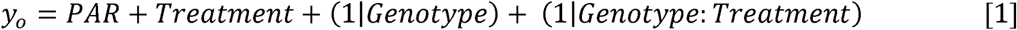

Since most of the datapoints were in the light limiting phase, resulting in a linear dependency on the light intensity, the exposed photosynthetic active radiation (PAR) and treatment were implemented as a stand-alone fixed effect, the genotype was defined as a random effect in addition to the interaction of genotype to treatment, defined as genotype-by-environment (GxE). The variance component of genotype as well as its interaction with the treatment, was used for statistical analysis and identifying photosynthetic QTL.

### Leaf lipidomics

Fully developed leaflets were harvested ten days post-stress treatment in the morning and flash-frozen under liquid nitrogen, followed by an extraction as described in Hummel, et al. ^72^ and modified according to Bulut, et al. ^73^. Briefly, a phase-separation based extraction method was performed using methyl *tert*-butyl ether:methanol (3:1 v/v) and water:methanol (3:1 v/v). The organic phase is used for lipid analysis. Dried aliquots were resuspended in acetonitrile:isopropanol (7:3 v/v) for lipids. For lipid annotations, an in-house library was used. In addition, each set of samples run on the analytical platforms were guided by several equally distributed pooled quality controls to correct for chromatographic shift in retention time and signal intensities across the data acquisition.

Chromatogram processing was conducted using MS Refiner (Expressionist 12.0). This includes chromatogram alignment, peak detection, noise filtering and isotope clustering. In addition to the targeted approach, we considered identities for detected compounds from publicly available legume metabolomics and lipidomics resources ^74^.

### Calculation of best linear unbiased predictors (BLUPs), heritability and data transformation

To obtain BLUPs for each multiomic trait (y_mo_) a mixed linear model (LMM) was generated, with slight variation by the exclusion of PAR from equation 1:

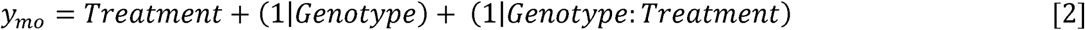

where treatment was implemented as a stand-alone fixed effect, the genotype as a random effect in addition to the interaction of genotype to treatment defined as GxE, similar to equation 1.

Further, broad sense heritability was calculated as follows:

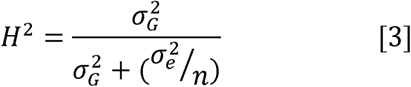

where H^2^ is broad-sense heritability, 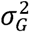 the genotypic variance, 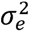 the error variance and *n* the number of independent biological replicates.

To achieve normal distribution, were transformed using Box-Cox power transformation ^75^:

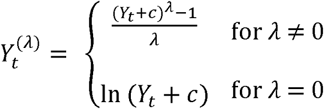

### Genome-wide association study

The polymorphic variants called from the whole-genome sequenced *P. vulgaris* population ^7^ were filtered for 5% minor allele frequency and 10% missing rate. Genome-wide association study was carried out using single-locus mixed linear model (MLM; Zhang, et al. ^76^), in which population stratification was fitted as a fixed effect, while the kinship ^77^ among individuals was incorporated using the variance-covariance structure of the random effect for the individual in the R package rMVP ^78^. The Bonferroni correction was carried out after determining the true number of independent markers by pruning SNPs by linkage decay using the R package SNPrelate ^79^ with the argument ‘100kb 10kb 0.2’ resulting in an adjusted p-value of 2.014e^-6^.

### Proteome files retrieval and inference of orthogroups, orthologs, and gene families

Proteome FASTA files of *Phaseolus vulgaris*, *Arabidopsis thaliana*, *Solanum lycopersicum*, *Oryza sativa*, *Zea mays*, *Glycine max* were retrieved from the Phytozome repositories. The orthogroups were inferred from protein FASTA files using OrthoFinder (Emms and Kelly, 2015; v2.2.7) as described in Naake, et al. ^80^.

### Biological Validation

Selected candidate genes were cloned under a constitutive promoter (35S) with C-terminal fusion of a green fluorescence protein (GFP) according to Zhang, et al. ^81^ to determine subcellular co-localization using transient overexpression as described in Zhang, et al. ^82^ modified for *Nicotiana benthaminana*. In brief, RNA from *P. vulgaris* was isolated using a NucleoSpin® RNA plant kit (Macherey-Nagel) according to the manufacturer’s instructions. First-strand cDNA was synthesized using 1.5 μg RNA and Prime Script™ RT reagent Kit with gDNA eraser (Takara) according to the manufacturer’s instructions. The entry clone was obtained through recombination of the PCR product with pDONR207 (Invitrogen). By LR recombination error-free clones were introduced into pK7FWG2. The transformation of leaves of *N. benthamiana* was performed following Karimi, et al. ^83^, and *Agrobacterium tumefaciens* (AGL1) containing vector pBin61-p19 was infiltrated with an OD600 0.5. DM6000B/SP5 confocal laser scanning microscope (Leica Microsystems, Wetzlar, Germany) was used for verification of expression. Lipidomic shifts were analyzed after 1 and 3 days using 50 mg of transiently transformed *N. benthamiana* leaves.

Further, overexpression constructs were used for hairy root transformation in *P. vulgaris* ^84^. After the stem was inoculation with *R. rhizogenes*, the infection site was kept at high humidity to allow hairy roots to develop. After sufficient hairy root biomass was generated, hairy roots were collected under liquid N for lipidomic characterization.

## Authors’ contribution

S.A. designed and conceptualized the experiments. A.R.F. and K.K. provided assistance. R.P., E.Bi. and E.Be. provided the genetic resource. Data generation was performed by M.B. and S.A.. M.B. processed and analyzed the data. M.B. and S.A. wrote the manuscript. All authors revised the manuscript.

## Acknowledgments

Thanks to all the people from the Max-Planck Institute of Molecular Plant Physiology who helped in either maintaining and harvesting the plants and providing the computational environment to process and analyze the multiomic data. M.B. acknowledges the funding by the IMPRS-MolPlant. R.P., A.R.F. and S.A. acknowledge the financial support for the EU Horizon 2020 research and innovation Programme, project INCREASE (grant agreement No. 862862). A.R.F. and S.A. acknowledge BG16RFPR002-1.014-0003-C01 project, financed by the European Regional Development Fund through the Bulgarian Program for Research, Innovation, and Digitalisation for Smart Transformation (PRIDST) Operational Programme. S.A. acknowledges the NATGENCROP project: HORIZON-WIDERA-2022- TALENTS-01, No. 101087091.

